# Longitudinal neural and vascular recovery following ultraflexible neural electrode implantation in aged mice

**DOI:** 10.1101/2022.08.08.503247

**Authors:** Fei He, Yingchu Sun, Yifu Jin, Rongkang Yin, Hanlin Zhu, Haad Rathore, Chong Xie, Lan Luan

## Abstract

Flexible neural electrodes improve the recording longevity and quality of individual neurons by promoting tissue-electrode integration. However, the intracortical implantation of flexible electrodes inevitably induces tissue damage. Understanding the longitudinal neural and vascular recovery following the intracortical implantation is critical for the ever-growing applications of flexible electrodes in both healthy and disordered brains. Aged animals are of particular interest because they play a key role in modeling neurological disorders, but their tissue-electrode interface remains mostly unstudied. Here we integrate in-vivo two-photon imaging and electrophysiological recording to determine the time-dependent neural and vascular dynamics after the implantation of ultraflexible neural electrodes in aged mice. We find heightened angiogenesis and vascular remodeling in the first two weeks after implantation, which coincides with the rapid increase in local field potentials and unit activities detected by electrophysiological recordings. Vascular remodeling in shallow cortical layers preceded that in deeper layers, which often lasted longer than the recovery of neural signals. By six weeks post-implantation vascular abnormalities had subsided, resulting in normal vasculature and microcirculation. Putative cell classification based on firing pattern and waveform shows similar recovery time courses in fast-spiking interneurons and pyramidal neurons. These results elucidate how structural damages and remodeling near implants affecting recording efficacy, and support the application of ultraflexible electrodes in aged animals at minimal perturbations to endogenous neurophysiology.

## 1 Introduction

Cortically implanted neural electrodes offer precise recordings from individual neurons (single-units)(*1*), several neurons (multi-units), and a small tissue volume (local field potentials (LFPs))(*2*). They have been an important neurotechnology for understanding the mechanisms, functions, and disorders of the nervous system, and for enabling brain-machine interface that restores or augments reliable communication between the brain and the external world (*3–5*). Rigid neural electrodes, such as Utah arrays(*6*), silicon microelectrodes(*7*), tetrodes(*8*) and microwire arrays(*9*), have contributed significantly for both basic neuroscience and translational applications. However, their intracortical implantation elicits a robust and well-characterized foreign body response that leads to progressive neurovascular damage over the implantation duration (*10, 11*), which hinders their chronic applications.

The needs to suppress the neuroinflammatory response and improve the tissue-electrode interface have fueled the development of flexible neural electrodes in the past decade (*5, 12, 13*). Particularly, we (*14–16*) and others (*17–20*) have shown that ultraflexible electrodes of a thickness of about 1 μm elicit minimal foreign body responses during chronic implantation of several months. Post-mortem histology examined the distribution of key cell types such as neurons and astrocytes in hippocampus and cortex, showing a relatively uniform cell distribution without obvious depletion of neurons or enhancements of astrocytes near the implants at multiple chronic time points after implantation (*16–20*). Longitudinal two-photon (2P) imaging repeatedly mapped the tissue-electrode interface, which revealed vascular remodeling in the first several weeks and a relative static tissue-electrode interface onwards featuring intact blood-brain barrier, no chronical neuronal loss, and no astrocytes accumulation (*14, 15*). These studies have provided convincing evidence that the ultraflexible electrodes significantly reduce long-term tissue invasiveness compared with their rigid counterparts in young adult animals.

Implantation of ultraflexible electrodes often requires a temporary attachment to a rigid shuttle device(*21–23*). The penetration of the shuttle-electrode dual through the neocortex inevitably exerts mechanical pressure, raptures the vascular beds around the insertion sites, and creates a stab wound that has a larger footprint than the footprint of the electrode itself. Emerging evidence suggest that microscopic injuries in the vasculature could have long-lasting, spatially spread-out impairment of neural activity(*24*). Therefore, the structural and functional responses of local vascular beds to the implantation trauma and its correlation with neuronal activity is critical to understand the tissue-flexible-electrode interface and how brain responses to slab injury. However, this is only scarcely and insufficiently studied. A significant limitation is that immunohistochemical techniques, the most commonly used methods to examine tissue-electrode interface, are ill-suited to probe neuronal and vascular dynamics because they require all analysis to be conducted post-mortem at one time point.

Another crucial but commonly overlooked factor is the animal’s age. Aged animals have different metabolism and immune responses than the young, adult counterparts(*25, 26*). Therefore, they more accurately model the aged human population of which neurological disorders become prevalent(*27*). But for the same reasons their neurovascular responses to implantation damage could be distinct from young adult animals, particularly given the documented attenuation of brain response to vascular endothelial growth factor-mediated angiogenesis and neurogenesis(*28*). In stark contrast to the growing awareness of the necessity of aged animals in studying neurological disorders and aging-related neuropathology, no study to date has examined their tissue-electrode interface, which unavoidably casts doubt on the application of these novel electrodes in the neuropathological studies in aged animal models.

Our goal is to better understand the temporal evolution of vasculature surrounding ultraflexible electrodes and how vascular remodeling correlate with neuronal activity while the brain recovers from the implantation damage in aged mice. Built upon our previous development of ultraflexible NanoElectronic Threads (NETs) (*14, 15*), we engineered and optimized the fabrication materials and procedures for a transparent, photo-inert substrate to support repeated *in vivo* 2P imaging without confounding phototoxic effects. We performed longitudinal *in vivo* 2P imaging of vasculature and microcirculation, and carried out electrophysiological recordings consecutively for six weeks following the implantation of ultraflexible NETs. We chose to study mice at 17 months and older, for which the aging effects on cortical reorganization have been demonstrated (*29, 30*).

## 2 Materials and Methods

### 2.1 Ultraflexible NET fabrication and preparation

To optimize for concurrent, longitudinally *in vivo* 2P imaging at the tissue-electrode interface, we made two modifications on the materials and fabrications of NETs. First, instead of using photoresist SU-8 as the backbone material, we used polyimide (PI) (PI2574, HD Microsystems, NJ, USA) to construct the insulating layers. PI offers improved tensile strength and biocompatibility than SU-8 while maintaining a similar level of fabrication convenience. Second, to suppress the photovoltaic effect, we used insulating glass substrate, instead of silicon. The total thickness of PI-NETs fabricated in this study is about 1 μm, similar as their SU-8 counterparts. The key steps of fabrication procedure are illustrated in Fig. 1A and outlined as follows. (*1*) An 80 nm-thick nickel sacrificial layer was patterned using photolithography and electron beam evaporation, which served as the final release layer for the ultraflexible section of NETs. (*2*) PI thin film was spin-coated at a thickness of 450 – 550 nm and thermally treated. (*3*) Trace-lines connecting the bonding pads with individual recording sites were patterned using photolithography and electron beam evaporation of Cr/Au at a thickness of 5/120 nm. Trace-lines were 1.5 μm in width and 3 μm in pitch in the implanted section. (*4*) Bonding pads were defined using photolithography and metal deposition of Cr/Ni/Au at a thickness of 5/80/20 nm. (*5*) A second layer of PI was spin-coated and thermally treated. An etching mask was patterned using photolithography and reactive ion etching (RIE) by O_2_ plasma was used to patten via and define the mechanical structure of NETs. After etching, the total thickness of PI shanks was at 900 – 1100 nm. (*6*) Individual recording sites in the implanted section were patterned by photolithography and metal deposition using Cr/Au at a thickness of 5/120 nm. Two designs of 4-shank NETs were used in this study. The first design, used in Fig. 1 – 3 and Fig. 5 & 6, had 32 channels in total, 8 recording sites per shank at a contact spacing of 100 μm center-to-center and a contact size of 30 μm in diameter. The shank width was 50 μm. The second design, used in Fig. 5, had 128 channels in total, 32 recording sites per shank at a contact spacing of 25 μm center-to-center and a contact size of 25 μm in square. The shank width was 96 μm. The inter-shank spacing when implanted was about 400 – 500 μm.

**Fig. 1:**
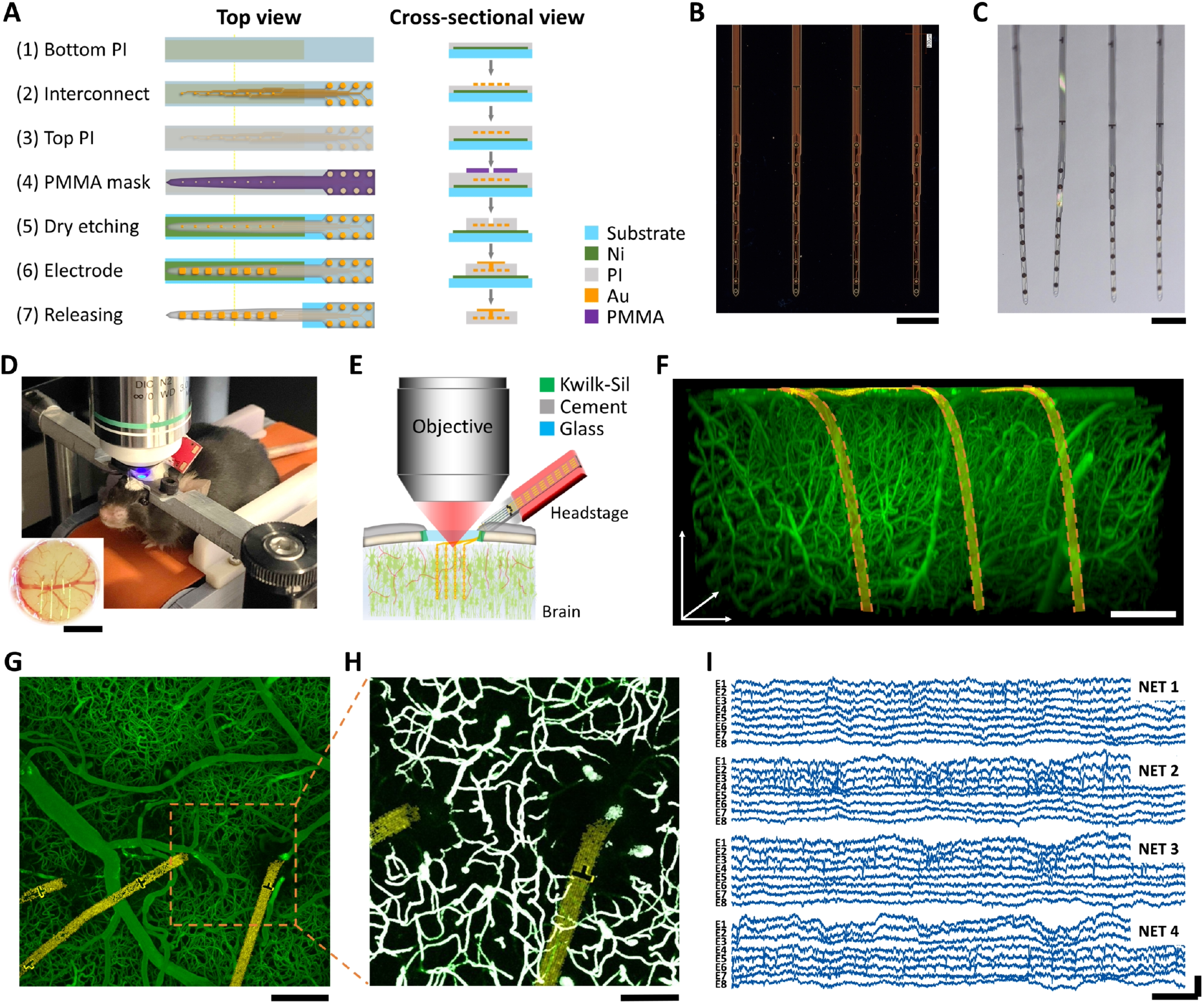
Ultraflexible PI NET permits longitudinal 2P imaging of microvasculature and concurrent neural recording. **A:** key microfabrication steps of PI-NET. **B, C:** photos of NETs (4 shank, 32 channel in total) on the fabrication substrate (B) and after released from the substrate and immersed in water **(C)**. **D, E:** Photo **(D)** and schematics **(E)** showing NET coimplanted with a cranial window to accommodate concurrent 2P imaging and electrophysiological recordings longitudinally. Animal was awake and head-fixed for all measurements. **F:** a representative 3D reconstructed image of vasculature surrounding three NET shanks at 6 weeks after implantation. Golden ribbon: pseudo-colored NETs. **G:** a representative maximum intensity projection (MIP) in xy plane (at 200 μm deep). **H:** A zoomin binarized image of G for quantification of micro-vessel volume fraction. White: binarized result. **I:** representative laminar cortical recordings from a 32-channel NET array at week six after implantation. Scale bars: 200 μm **(B, C, F, G)**, 100 μm **(H)**, 1 mm (inset in **D**); 20 ms (**I**, horizontal) and 500 μV (**I**, vertical).

After fabrication, a customized print circuit board (PCB), the detail of which was previously reported (*31, 32*), was mounted on the matching contact pads on the glass substrate. The implantable section of the probe was then soaked in nickel etchant (Type I, Transene Inc., MA, USA) for 5 mins at the room temperature to release the ultraflexible section of the probe, whereas the contact region remained attached to the substrate. The substrate was cleaved in water to the desired length before implantation. The contacts were electrochemically deposited with PEDOT:PSS to lower the impedance to ~30 kΩ at 1 kHz in physiological saline. The released sections of NETs were attached onto shuttle devices made of tungsten microwires of dimeters at 50 μm (W5574 Tungsten Wire, Advent Research Materials, Oxford, England) using bio-dissolvable adhesive 2.5% (m/v) PEG (PEG 35k, Fisher Scientific, PA, USA) solution. The NET-shuttle device assembly was sterilized in an ethylene oxide sterilizer (Anprolene AN74, Andersen Products Inc., USA) before implantation.

### 2.2 Animals and Surgery

A total of 12 aged male C57BL/6J wild type mice (National Institute on aging Aged Rodent Colonies) were used in the experiments. The animals’ age was 16 – 17 months at the NET implantation and was 18 – 19 months at the end of measurement period. They were housed individually after NET implantation at a 12-hr, 7:00 to 19:00, light-to-dark cycle. All animals received standardized cage supplementation (cage enclosures, nesting material and objects to gnaw) with water/food *ad libitum.* All animals received one surgery during which a NET electrode array was implanted intracortically and a chronic cranial window was mounted on the skull(*31*).

Mice were anesthetized with medical O_2_ vaporized isoflurane (3%) in an induction chamber and then placed prone in a stereotaxic frame (David Kopf Instruments, California, USA) via nose-cone inhalation of medical O_2_ vaporized isoflurane at 1.5 – 2% (Kent Scientific, Connecticut, USA). Ethiqa XR (3.25 mg/kg) and dexamethasone (2 mg/kg) were administrated subcutaneously to reduce pain and inflammation during the craniotomy and implantation procedure. Body temperature was maintained at 37 °C with a feedback heat pad (Kent Scientific, Connecticut, USA). Heart rate and breathing rate were monitored by the surgeon. A circular portion of skull (3 mm in diameter) atop the motor and somatosensory cortex was removed with a dental drill (Ideal Microdrill, 0.5 mm burr, Fine Science Tools, CA, USA) under constant sterile artificial cerebrospinal fluid (buffered pH 7.4) flushing. Dura matter was removed to facilitate a larger imaging depth in the following imaging sessions. 4-shank NET 2D arrays were implanted stereotaxically using tungsten microwires as the shuttle device and bio-dissolvable adhesive as discussed in previous publication (*24, 31, 33*). The implantation angle of NETs was typically perpendicular to the brain surface. After implantation, the carrier chip was carefully positioned and mounted on the remaining skull at a 55-degree angle and about 5 mm away from the implantation sites to accommodate the objective lens of the 2P microscope (Fig. 1D, E). A small hole was drilled on the contralateral hemisphere of the brain and a stainless-steel wire was inserted into the brain as the grounding reference for later electrical recording. A 3 mm round cover glass (#1, World Precision Instruments, Sarasota, FL, USA) was placed over the exposed, NET implanted brain area with a layer of sterile saline between the two. Gentle pressure was applied to the cover glass while the space between the coverslip and the remaining skull was filled with Kwik-sil adhesive (World Precision Instruments, FL, USA). A layer of vet-bond tissue adhesive (3M, USA) was applied to cover the skull and the Kwik-sil. Then an initial layer of C&B-Metabond (Parkell Inc., NY, USA) was applied over the cyanoacrylate and the vet-bond. This process ensured a sterile, air-tight seal around the craniotomy and allowed for restoration of intracranial pressure. A second layer of Metabond was used to cement the coverslip and the NET carrier chip to the skull. A final layer of Metabond was used to cement a customized titanium headplate for later head-constrained measurements. This surgical preparation allows longitudinal 2P imaging at the NET implantation locations and concurrent electrophysiological recordings so that we can evaluate the neural and vascular dynamics after NET implantation.

All experiments were approved by the Institutional Animal Care and Use Committee (IACUC) at Rice University and comply with the National Institutes of Health guide for the care and use of Laboratory animals.

### 2.3 2P imaging

2Pimaging was performed using a multiphoton laser scanning microscope (Ultima 2P Plus, Bruker Corporation, MA, USA) equipped with a 16× water immersion objective (NA = 0.8, Nikon, Japan). A widely tunable ultrafast laser (InSight X3, spectra-physics, CA, USA) was tuned to 920 – 930 nm to provided 2P excitation at 80 MHz repetition rate. A field of view (FOV) of 1.1 mm × 1.1 mm, 1024 × 1024 pixels, dwell time of 2 – 3 μs were typically used to acquire vasculature images. Image stacks were collected at 1 – 2 fps from the brain surface to a final depth at which signals could no longer be detected (typically 500 to 600 μm), with a 2-μm Z-step size. Laser power was increased logarithmically during depth stepping to account for exponential light penetration losses within tissue. Fluorescence emissions were detected simultaneously by two GaAsP photomultiplier tubes (Hamamatsu, Japan) with a 595/50-nm filter (Semrock) for “red” fluorescence emission and a 525/70-nm filter (Semrock) for “green” fluorescence emission. Mice were unanesthetized, head-fixed on a free-moving treadmill during imaging. Prior to imaging, mice were anesthetized and administered fluorescein isothiocyanate (FITC)–dextran (0.1 ml, 2 M MW, 5% w/v, Sigma-Aldrich) via tail veil to label blood vessels. NET was not fluorescently labeled and was detected by the autofluorescence from the metallic lines and markers.

To measure the red blood cell (RBC) speed in the capillaries, a capillary of interest was identified after obtaining a 2P image of vasculature at the desired depth, and the imaging frame was electromechanically oriented to the direction of the scan along it. Repetitive line-scans were performed along the central axis of a capillary to detect the motion of unlabeled RBCs, which appeared as dark stripes in these line-scan measurements (*34*). A distance of 20 –40 μm with 128 pixels per line were scanned along the capillary, repeating 128 times at 1 to 2 ms per scan.

### 2.4 Vascular Analysis

Fiji (Open source, ImageJ.net) and Matlab (Mathworks, MA, USA) was used to preprocess 2P images of vasculature. The pre-processing has two steps to achieve homogenous intensity across all three dimensions in each imaging stack and across all imaging stacks of each animal (Supplementary Figure SFig.1). First, we corrected the variations in illumination along depth and across multiple longitudinal sessions. The acquired images were smoothed by a 3D Gaussian filter with a 1.09 μm full width at half maximum (FWHM) kernel size for background blurring and homogenization of vessels. Then background subtraction using a rolling ball algorithm was conducted to enhance the signal-to-noise ratio (SNR)(*35*). After smoothing and background subtraction, all longitudinal imaging sessions of the same animal were grouped together to compute the grand mean and the standard deviation (SD) values of all image sessions. Z-slice normalization was performed to adjust the mean and SD values of each individual Z-slice images to the grand mean and SD values of all images to correct the illumination attenuation from physical depths and compensate for different exposure conditions at each session.

Second, we corrected non-homogenous illumination of the *xy* plane, resulting from a gradient of tissue depth, shadowing from surface vessels, or spotty window clouding. We generated maximum intensity projection (MIP) image excluding 100 μm of apical surface, and obtained an intensity bias field by applying 2D Gaussian filters of three different FWHM kernel sizes on the MIP image. The Gaussian kernel sizes were typically 8 μm, 14 μm, and 30 μm respectively to count for non-homogenous intensity with different spatial profiles. The intensity was then corrected using:

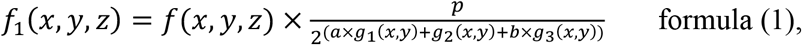

where *g_i_*(*x,y*) was the intensity bias field at the three kernel sizes; *f*(*x,y,z*) was any individual image in the Z-stack after across-session Z-normalization; *a, b, p* were constants adjusted for each mouse to achieve homogenous intensity in the *xy* plane. Typical values of *a* and *b* were [0.5 1]; and *p* were [10 30]. These procedures produce relative uniform intensity and contrast throughout a FOV of 1.1 mm × 1.1 mm × 0.5 mm as shown in Fig. 1F. Pre-processed imaging stacks were then sectioned in different dimensions and projected in a MIP image. A representative MIP in *xy* plane is shown in Fig. 1G. The pre-processed image stacks are also sectioned and projected into MIP images in *yz and xz* plane*s* at various thicknesses to visualize the vasculature-NET interface. Sessions and implantation locations that have severe shadowing effects from bleeding, thickening of dura mater, and overflow of cement when mounting the cranial window were excluded from the analysis.

After pre-processing, we use the “local contrast” method in the IMARIS software (Bitplane, Belfast, United Kingdom) to generate binary imaging stacks (Fig. 1H). This method overestimates the vessel diameters because the contrast threshold for binarization is typically lower than the contrast at the FWHM of the pre-processed images. To obtain the vessel volume fraction, we corrected the binarized volume fraction by a scaling factor to compensate the overestimation. The scaling factor was determined by measuring the FWHM of vessels in the pre-processed maximum projection, computing its ratio to the width of the same vessel after binarization, and averaging among all randomly selected vessels (120 vessels per animal).

To quantify the vessel-density change in three dimensions across time, we further converted the binarization images into a volume fraction map by dividing the FOV into voxels of 30 μm × 30 μm × 50 μm and computing the volume fraction of vessels in each voxel. For a typical FOV of 1120 μm × 1120 μm × 500 μm, we obtained a 38 × 38 × 10 matrix representing the local volume fraction of the vessels per voxel. Then we generated the heatmap of the recovery speed around the NETs by averaging three voxels along Y axis (90 μm thickness in total), which was defined as the time needed to reach 80% vessel density relative to the unaffected regions at the same cortical depth.

We also used FIJI to get the skeleton and diameter map of the vascular for further analysis of the vascular morphology. We first generated the skeletons of the vascular image by using the 3D skeletonize plugin in Fiji and merged the diameter map together to classify vessel segments into large vessels (>14 μm), medium vessels (≼14 μm), and capillaries (≼ 9 μm). To extract the diameter of vessels, the vessel was skeletonized in the MIP images in the *xy* plane. The intensity profile of a line perpendicular to the vessel axis was taken at every pixel along the skeleton, averaged across a running window of five pixels. The diameter was calculated from the FWHM of this intensity profile using the Local Thickness plugin in Fiji. To counter the problem of large surface vessels being classified as two or more small vessels due to significant variations in diameters along the vessels, we used the VasoMetrics plugin(*36*) in Fiji to manually select the central line of the vessels and then quantified the diameter of the vessels as the FWHM of the intensity profile perpendicular to the central line.

### 2.4 Quantification of capillary blood flow

To quantify the RBC velocity, flux, and hematocrit values from 2P line scans along the axis of capillaries, line-scan images were cropped in ImageJ to include regions with a high SNR. An open-source MATLAB script from Drew *et al* (*37*) was adapted to calculate the RBC velocity using Radon transform. A customized MATLAB script was used to count the number of RBC streaks to determine the RBC flux. To calculate hematocrit values within a capillary over time, line-scan frames were preprocessed and binarized using methods outlined in (*38*), and the percentage of dark stripes (*i.e.,* containing RBCs) was calculated within each frame period (3 – 4 ms). The noisy images after binarization steps were excluded from the study and therefore 466 capillaries in total were included in the hematocrit statistics plot. The accuracy of the automated scripts to quantify RBC velocity, flux, and hematocrit values were verified by manual calculation of a subset of the data.

### 2.5 Electrophysiology Data Acquisition and analysis

To evaluate NET performance and detect neuronal electrical activities over time, voltage signals from intracortically implanted NET contacts were acquired from awake, head-fixed mice on a free-moving treadmill using a 32-channel RHD 2132 acquisition system (Intan Technologies, CA, USA). The sampling frequency was 30 kHz, and a band pass filter at 0.5 Hz – 6000 Hz was applied. Representative recoding traces were shown in Fig. 1I. Impedance of all contacts was measured before each session, and contacts with impedance at 1 kHz larger than 2 MΩ were regarded as bad connections and excluded from the measurements. Each recording session lasted about 30 min.

Spectrograms of LFPs were computed by down sampling the neural recording data to 1 kHz and performing Fourier transform using a 1-s non-overlapping window. Integrated power in different frequency bands, *i.e.,* 30 – 60 Hz and 60 – 110 Hz was calculated to extract the envelope of the signal. Multi-unit activity (MUA) was obtained by applying a bandpass filter at 300 – 3000 Hz. For longitudinal comparisons, a recording duration of 30 mins excluding sections with heightened motion artifacts was used to compute the mean and SD, and all values are normalized to the mean of the last day of measurements (six weeks after implantation).

Spike sorting was performed by thresholds detection and clustering in MountainSort (*39*). Adjacency radius was set to 100 μm to restrain the clustering neighborhoods to the immediate neighboring recording electrodes. The event detection threshold was set to 3.5 SD. Manual curation of sorted units was then performed to reject noise clusters and combine over-split clusters. Units are classified as single unit recordings if ≤ 1% of the inter-spike intervals (ISIs) are below 2 ms. The spike-waveform width (trough-to-peak) was assessed based on the duration from the troughs to the rising peaks on the waveform. Autocorrelograms were fitted with a triple-exponential equation to determine the temporal scale of the rising phase of the ACGs (*τ*_rising_) as defined in reference (*40*). Cells are classified into two putative cell types: narrow interneuron are assigned if trough-to-peak width ≤ 0.425 ms, while the others are remaining putative pyramidal cells.

### 2.6 Statistics

The sample size was determined by power analysis using G*Power. The blood flow speed, flux, and hematocrit were statistically evaluated by one-way ANOVA with Tukey’s *post hoc* test for multiple comparisons.

## 3 Results

We used a suite of longitudinal 2P imaging and electrophysiological recording methods to examine and longitudinally track the vascular structural remodeling and functional recovery, neural activity, and their correlation with vascular recovery in aged mice. These longitudinal measurements started immediately after the surgical implantation of NETs and lasted till six weeks post implantation after we had observed lasting stability. All measurements were performed on unanesthetized, head-fixed animals to remove confounding effects from anesthesia.

### 3.1 vascular remodeling and neural recovery were the most pronounced in the first two weeks of implantation

We first studied the longitudinal structural remodeling of the vasculature surrounding the implantation sites of 4-shank, 32 channel NETs in aged mice. We used tungsten wires of 50 μm in diameter as shuttle devices to implantation NETs. Therefore, we expected a wound of similar or larger size spanning the cortical depth to appear immediately after implantation and evolve over time. Fig. 2A shows representative time series of vasculature-NET interface from the implantation day to six weeks after, for which *xz* plane MIPs centered at the implantation site depicted the vasculature evolution across the cortical depths. A range of thickness along the third dimension (Y axis) for MIP were tested and 90 μm was chosen to optimize visualization of the microscopic injury and its recovery over time: thinner projection along Y failed to include a completely volume of the injury around the implantation site, while thicker stacks included unaffected vasculature that veiled the damage (Supplementary Figure SFig.2). In this representative animal, the implantation induced local bleeding and raptures of vessels, leading to a lack of detectable vasculature in the close vicinity of NETs immediately after surgery (Day 0). Substantial vascular remodeling and angiogenesis had occurred by Day 7 and continued thereafter, resulting in a steady increase of vasculature over time that were the most pronounced in the first two weeks. Particularly, nearby penetration vessels that shuttle blood between the cortical surface and the parenchyma(*41*) reperfused by Day 7. By Day 28 the vascular remodeling had subsided and the vasculature surrounding NETs remained stable till Day 42, the last time point for the longitudinal measurements.

**Fig. 2:**
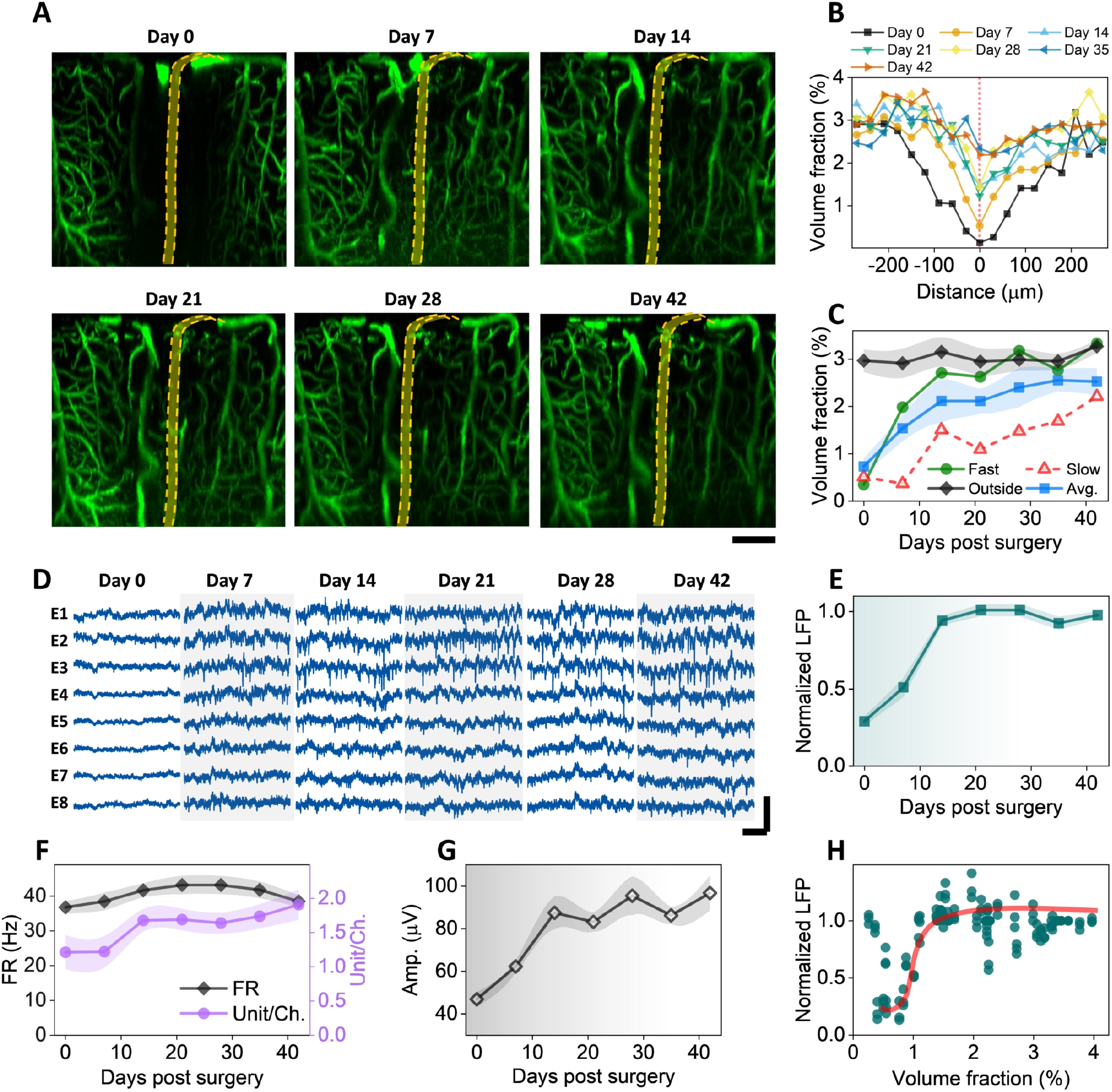
Vascular remodeling and neural recovery is the most pronounced in the first two weeks of implantation. **A:** MIP of *xz* sections (90 mm thick in Y) of a representative animal showing vascular recovery at the NET interface. Dashed orange lines depict NET. **B:** Volume fraction of vasculature as a function of distance from the NET showing the spatial extent of injury at multiple time points after implantation. **C:** Volume fraction of vasculature as a function of time post implantation from within 200 μm of the NET implantation site and more than 300 μm away; values are mean±SD (*n* = 16). **D:** representative laminar recordings from one NET shank (8 channels) showing spontaneous LFP and spiking activities. **E – G**: normalized LFP at 30 – 60 Hz **(E)**, firing rate (FR) and unit per channel **(F)**, and spike amplitude (G) as a function of days post implantation; values are mean ± SE (*n* = 30). **H:** scatter plot of normalized LFP and vasculature volume fraction showing a sharp increase in LFP at relative low vascularization. The solid line is guide for the eye. Scale bars: 100 μm **(A)**; 20 ms (**D**: horizontal), and 200 μV (**D**, vertical).

There were significant variations in the initial implantation trauma among animals. Supplementary Figure SFig.3 shows the time series of *xz* plane MIPs from another animal that had more severe initial damage than that shown in Fig. 2A. Deformation of brain tissue, a ‘dint’, occurred immediately after implantation, deteriorated at Day 7, and alleviated later. By Day 28 most vasculature around NET had re-perfused and tissue deformation was no longer detected. Summarizing data from *n* = 6 animals and *n* = 16 implantation sites, the volume fraction of vasculature as a function of distance from the NET shows that the spatial extent of the vascular injury monotonically decreased with time and the largest changes occurred in the first two weeks (Fig. 2B). Consistently, the volume fraction of vasculature within 200 μm of the NET implantation site monotonically increased with time after implantation (Fig. 2C). Variations among animals manifested as distinct recovery trajectories, while most significant increase of vasculature occurred in the first two weeks in all animals. As a control, the volume fraction of vasculature far away (> 300 μm) remained stable during the entire measurement periods at a value close to the recovered implantation site (Fig. 2C). Does the change in neural activity track the vascular recovery after implantation? Fig. 2D shows representative laminar recordings from one NET shank at eight cortical depths as days after implantation, from which LFP spectral power (Fig. 2E), spike rate, unit yield per channel (Fig. 2F), and spike amplitude (Fig. 2G) were quantified (methods). Here the spikes includes both single- and multi-units, and we will discuss single-unit recordings later. All these matrices suggest that neural electrophysiological activity increased in the first two weeks and remained stable afterwards. Notably, the relationship between LFP and vasculature volume fractions is nonlinear: LFP has a sharp increase at a relatively low volume fraction of vasculature (about 1%) and stayed relatively unchanged with increasing level of vascularization (Fig. 2H). This indicates that regeneration and reperfusion of a fraction of vasculature after implantation is sufficient to support the energy demands of neural activity at a population close to later periods. Further vascularization has little influence on the population-average neural electrophysiological activities.

### 3.2 Microvascular remodeling shows location and depth dependence, but these moderate differences do not affect electrophysiological recordings

We next examine if microvascular remodeling has any cortical depth or location dependence. Representative vasculature images at multiple cortical depths and time points after implantation (Fig. 3A) show two interesting features. First, vessels at shallower layers, including both pial vessels and micro-vessels, completed remodeling and reperfusion sooner than deeper layer microvessels. We analyzed vessel volume fraction in a cortical-depth specific manner and quantified the averaged time needed for regions within 200 μm of NETs to reach 80% vessel density relative to regions unaffected by NET implantations (Fig. 3B). While on average it only needed 10 days or less for the shallowest tissue (<200 μm of brain surface) to regain perfused vessels, the time needed for vessel recovery increased with cortical depth, and reached 30 days or more for regions deeper than 400 μm for the same level of vascularization. To map the spatial profile of vessel recovery with time, we quantified the vessels density in voxels and generate a heatmap (Methods), which depicted the time needed to reach 80% vessel density relative to the unaffected regions as a function of cortical depth and distance from the NET implantation sites (Fig. 3C). Consistently, deeper regions had a longer lasting, more spatially extended vascular deficits comparing with the shallower cortical regions.

**Fig. 3:**
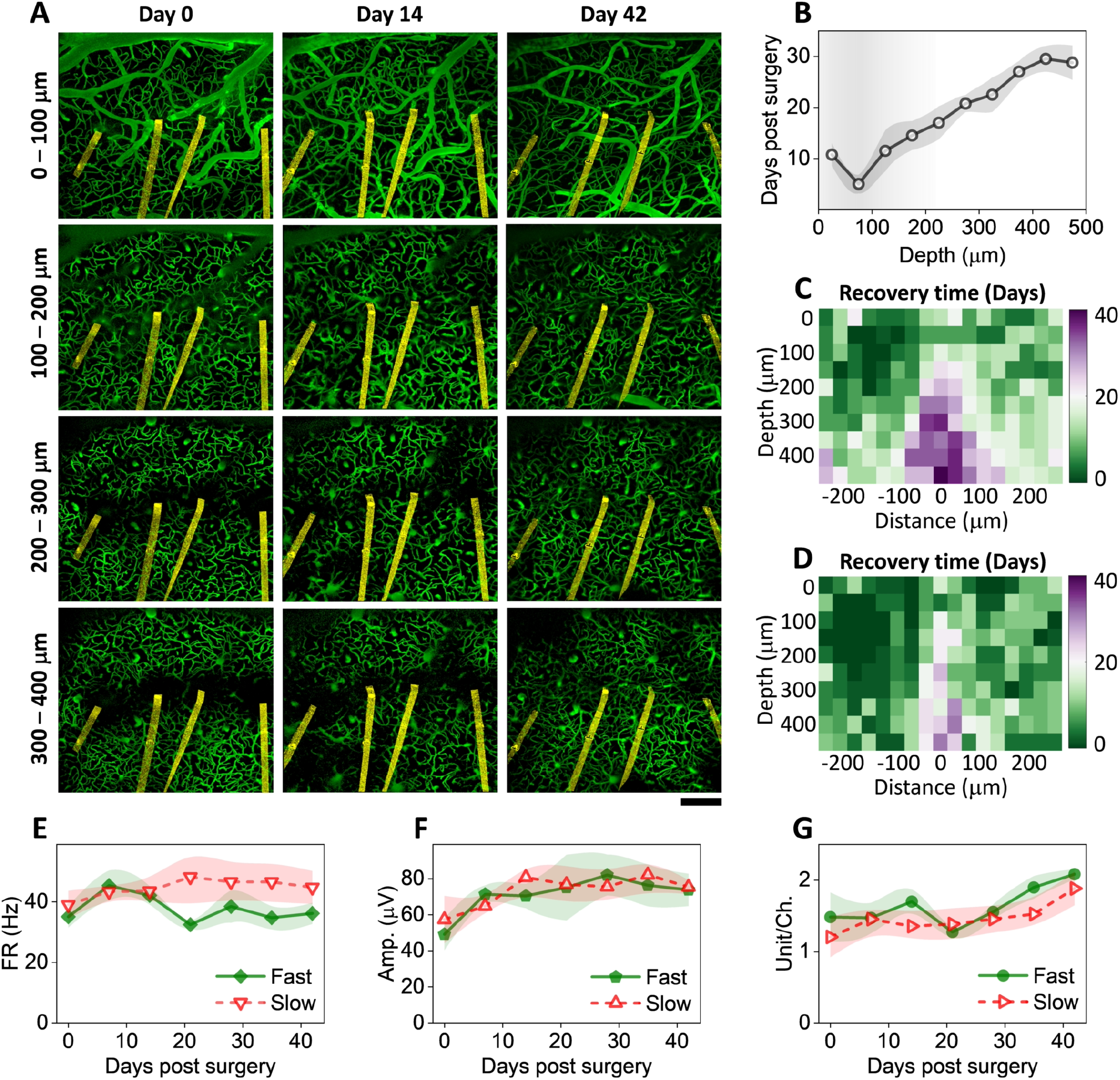
Microvascular remodeling shows location and depth dependence with little influence on neural electrophysiological activity. **A:** vasculature images at multiple cortical depths and time points after implantation from one representative animal. Arrow denotes the implantation site with faster vascular recovery. NETs are pseudo-colored as yellow. Scalebar: 200 μm. **B:** cortical-depth dependence of the averaged time needed for regions within 200 μm of NETs to reach 80% vessel density relative to regions far away. values are mean±SE (*n* = 12). **C:** color-coded vessel recovery time as a function of cortical depth and distance away from NETs (*n* = 12). **D:** color-coded vessel recovery time from a subset of animals in **C** representing faster-than-average recovery time course (*n* = 6). **E – G:** Spike rate **(E)**, amplitude **(F)**, and spike yield per channel **(G)** in the group of faster recovery vasculatures and the rest. No significant difference was detected; values are mean±SE (*n* = 9)

Second, vascular recovery at some locations is more prompt than others in the same animal (SFig. 4). In SFig. 4, the implantation sites away from major pial vessels (denoted by the arrow) led to faster vascular recovery. Vasculature recovery heatmap from the subset of implantation sites (Fig. 3D) that we designated as ‘faster recovery” showed less time needed for re-vascularization and reduced spatial extent of vascular damage in deeper regions comparing with the averaged cases shown Fig. 3C.

It is worth noting that these difference in vascular recovery did not influence the time course or magnitude involved in neural electrophysiological activity recorded by NETs. Fig. 3E – G show the spike firing rate (FR), the average amplitude of the spikes, and spike yield per channel in two groups: the fast recovery of vasculature as shown in Fig. 3D, and the rest animals. There is no detectable difference in neither of these electrophysiological matrices between the two groups. This is consistent with Fig. 2H. Because further vascularization exceeding a certain fraction of the normal vessel density has little impact on the population-average neural electrophysiological activities, the moderate difference in depth and location found in vasculature remodeling does not lead to significant difference in neural recording.

### 3.3 Large-scale single-unit recordings indicate similar time dependence of electrophysiological activity across cortical depths and cell types

To detect how individual neuron’s electrophysiological activity change after implantation in greater details, we implanted 4-shank, 128-channel NETs in another group of aged animals (*n* = 4) and performed more frequent measurements in the first two weeks post implantation. We employed a design of high recording site density, which allows for simultaneous detection of individual neurons by multiple channels (Fig. 4A), and therefore perform better in single unit isolation. Fig. 4B, C show the single unit yield per channel, the averaged firing rate (FR) of single units, and the average amplitudes of single units as a function of time. Notably, we detected a relatively large number of single units with relatively high spike rate on Day 0 comparing with the previous animal group (Fig. 2). We attribute the better recording on Day 0 to our deliberate delay of the first measurement to 8 – 12 hours post implantation in effort of removing possible influence of residence anesthesia from surgery. The spike amplitude increased monotonically with time in the first two weeks after implantation, which is consistent with the average distance between neurons and NET being reduced as the tissue-probe interface got tighter. On the other hand, the number of units and spike rate decreased significantly in the next few days, bounced back and surpassed Day 0 values by Day 10, and remained relatively stable onwards, suggesting a latency between the physical trauma and the affected neuronal activity. The most dynamic changes subsided by week 2, consistent with the time course of tissue-NET interface repair as indicated by longitudinal vascular imaging.

**Fig. 4:**
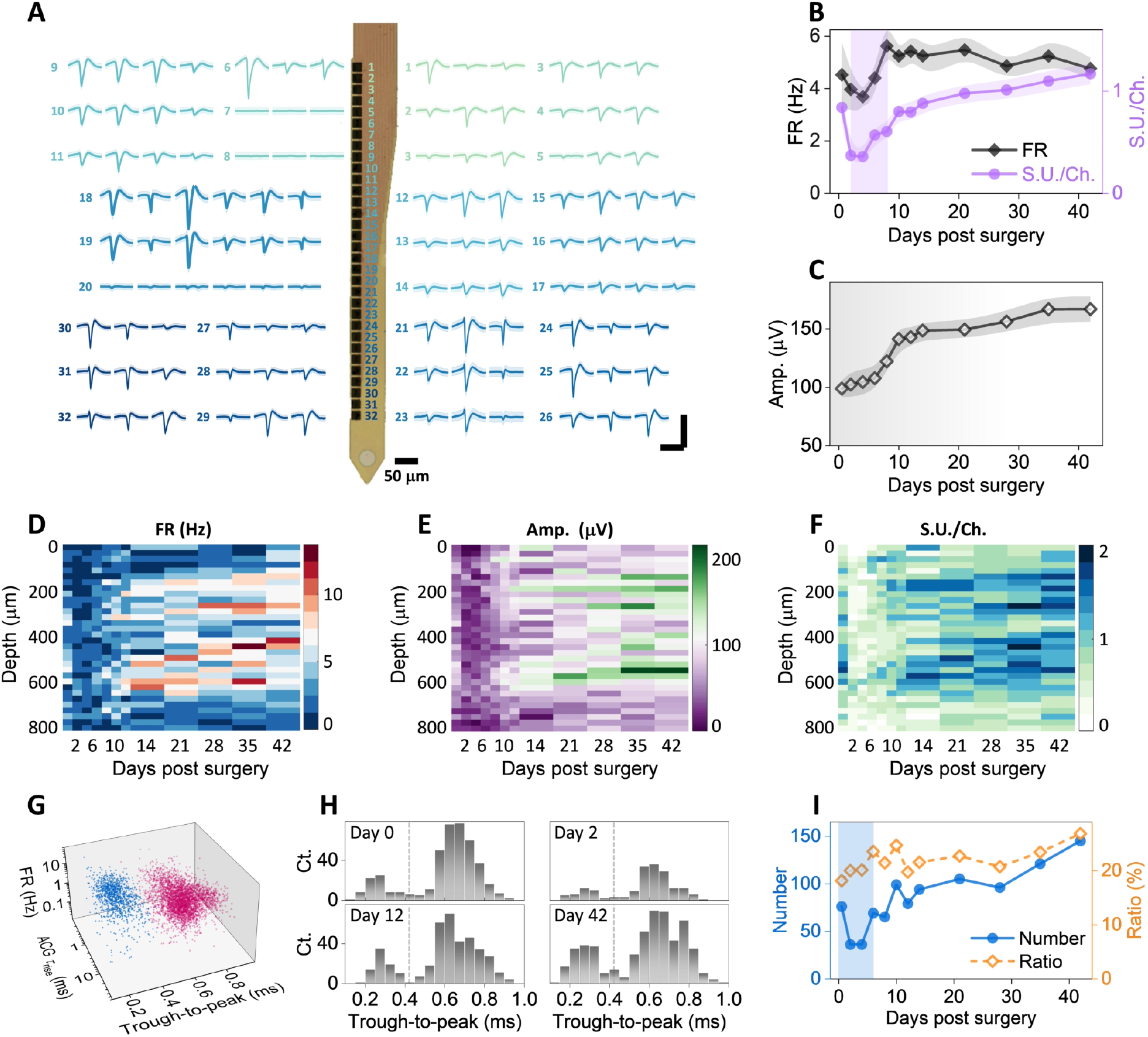
Large-scale recording reveals changes of single-unit activity over time, cortical depth, and cell types. **A**: Example recordings of single units from a single 32-channel NET shank. Numbers denote the recording sites along the shank. Multiple waveforms were detected at each site. Scale bars: 200 μV (vertical) and 2 ms (horizontal). **B – C**: single unit yield per channel, firing rate (**B**), and amplitudes (**C**) as a function of days after implantation; values are mean±SE (*n* = 16). **D – F**: cortical depth dependence of single unit firing rate **(D)**, amplitude **(E)**, and single-unit yield per channel **(F)** as a function of days post implantation. **G**: scatter plot of 4593 single units in 3D space for purtative cell classification (blue, interneurons; red, remaining neurons). **H:** example histogram distribution of trough-to-peak showing bi-modal distributions. **I:** Number of putative narrow waveform interneuron and their ratio as a function of days post implantation. Scale bars: 2 ms (**A**, horizontal), 200 μV (**A**, vertical).

Having a large number of individually addressed recording sites at a spacing of 25 μm and a coverage of 800 μm provides depth-resolved electrophysiological recordings as a function of time (Fig. 4D – F). Spike rate (Fig. 4D) and unit yield per channel (Fig. 4F) both show “bands” of more active regions at the depth of 200 – 350 μm and 450 – 650 μm. The location of these active regions is consistent with the anatomical locations of Layer 2/3 and Layer 4/5 neurons. Importantly, the two regions had similar level of electrophysiological activity at all time points measured, suggesting similar recovery time courses after implantation for Layer 2/3 and 4/5 neurons.

The ability to record a large number of single units also allowed us to putatively classify these neurons into different cell types and examine the electrophysiological activity of each type after implantation. We used the processing pipeline of CellExplorer (*40*) that classifies putative cell types based on three parameters, the width of the spike waveform (measured by the waveform “trough-to-peak”), the spiking frequency, and the burstiness of spiking (measured via the rise time of the autocorrelationgrams ACG *τ*_rise_). Fig. 4G shows a total of 4593 single units concatenating all measurement sessions, where two well-separated clusters were identified: narrow waveform (trough-to-peak ≤ 425 μs), fast-spiking interneurons, and the rest, mostly pyramidal cells. Furthermore, spike width histograms show bimodal distributions in all measurement sessions, including the most dynamic phases of the first 14 days and the relatively stable later stage (Fig. 4H), that quantitively agreed with the all-session clustering result. Importantly, the ratio of fastspiking interneurons remained stable at around 20% despite drastic changes in unit numbers in the first several days, suggesting that the implantation of NETs affects different cells similarly in this time window (Fig. 4I). This ratio of fast-spiking interneurons is similar to previously reported results at various time after implantation (*40*).

### 3.4 Vessel diameter and capillary blood flow remained relative stable with occasional vasospasm in the first two weeks

We next examined vessel diameters and capillary blood flow to detect if there were any vascular functional changes, and if so, when they occurred. To determine the vessel diameter changes over time, we classified vessels into three categories according to their diameters (Methods) and the functional classification of the vasculature tree: the large vessels represented pial vessels; medium-sized vessels were penetrating arterioles and venules; and small vessels were mostly microcapillaries (Fig. 5A). In these three vessel categories, the average diameters of medium and small vessels remained stable, while the large vessels on average showed a decrease over the time course of measurements (Fig. 5B).

**Fig. 5:**
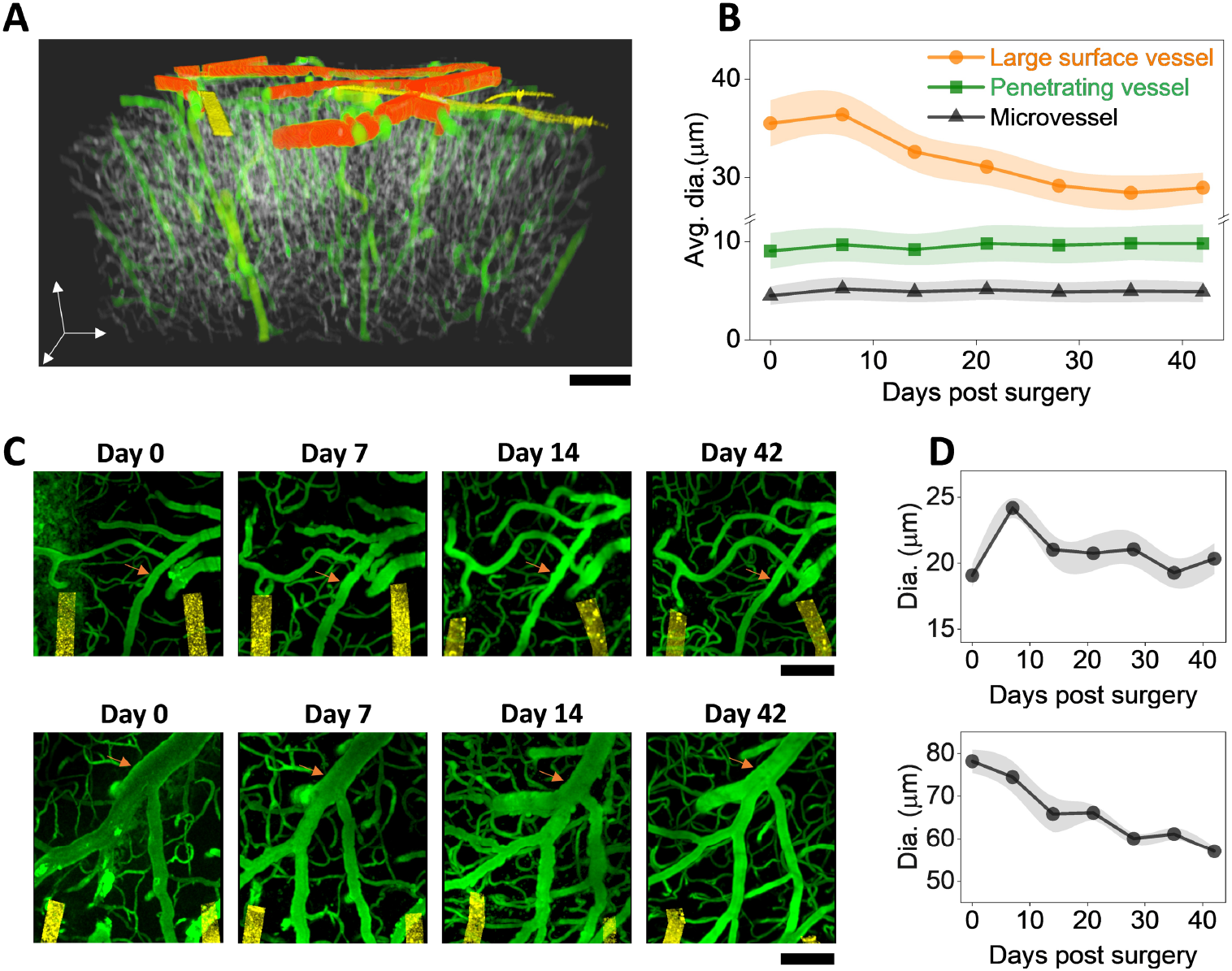
Vessel diameter remained relative stable but vasospasms were detected occasionally. **A:** color-coded binarized imaging stack showing the three categories of vessels. Orange: large vessels; green: medium vessels; grey: small vessels. Golden: NETs **B:** Vessel diameters as a function of time post implantation for the three vessel categories. **C:** 2P imaging stacks on the implantation day, Day 7, Day 14, and Day 42 post implantation from two animals showing diverse vessel changes. Vessel of interest marked in orange. All images are MIP of *xy* sections (100 μm thick stack in Z). **D:** Diameter of the marked vessel in **(C)** as a function of time post implantation; values are mean± SE (n = 7).

We then scrutinize the large vessels and discovered diverse changes with time post implantation. Notably, some of the major veins, particularly those close to the NET insertion sites, reduced in diameters significantly over time (Fig. 5C). While many other pial vessels had little changes in diameters over time, these “shrinking” veins led to the detectable reduction of vessel diameter when we averaged over all large vessels in all animals. Furthermore, we occasionally detected vasospasms sparsely distributed in the imaging FOV. The abnormalities in vessels were the most severe at Day 7, and subsided at Day 14 and after (Fig. 5D). We ruled out photodamage from 2P imaging as a possible cause because the compromised vessels were sparsely distributed in the FOV instead of concentrating to a subfield; and they were mostly penetrating vessels instead of the larger pial vessels on the surface which were more susceptible for photodamage. The averaged diameter of these compromised vessels increased at Day 7, and returned to a stable value one week later (Fig. 5D). The sparse vasospasms occurred when angiogenesis had picked up, but coincided temporally with the exacerbation of tissue deformation at Day 7 as we detected in some animals (discussed in SFig. 3), reflecting the multifaceted nature of vascular dynamics following injury. Multiple damaging and repairing signaling pathways are triggered by the implantation, and it is conceivable how they manifest in the angiogenesis and cerebrovascular complications depends on the timing and magnitudes of all factors involved.

Lastly, we measured microcirculation in the capillary beds to determine if microcirculation in perfused vessels changed over time after NET implantation. We scanned over 510 capillaries within 250 μm of NETs while staying away from damaged vessels, and quantified the RBC speed, flux, and hematocrit values at four time points from Day 7 to Day 42 post implantation (*n* = 6 animals). As angiogenesis and vascular remodeling took place with time, we included the perfused capillaries in the previous damage zone without distinguishing them from unaffected vessels in the vicinity (Fig. 6A, B). We sampled a similar density of capillaries per cortical depth from the brain surface to 500 – 600 μm into the cortex, and combined all measurements without considering the possible subtle difference at different cortical layers (*42*). RBC speed and flux at Day 7 were significantly higher than Day 14 but the effect size is small. Hematocrit remained stable over the six weeks recovery period (Fig. 6C-E). These results suggest that vascular structural remodeling has limited impact on the capillary microcirculation.

**Fig. 6:**
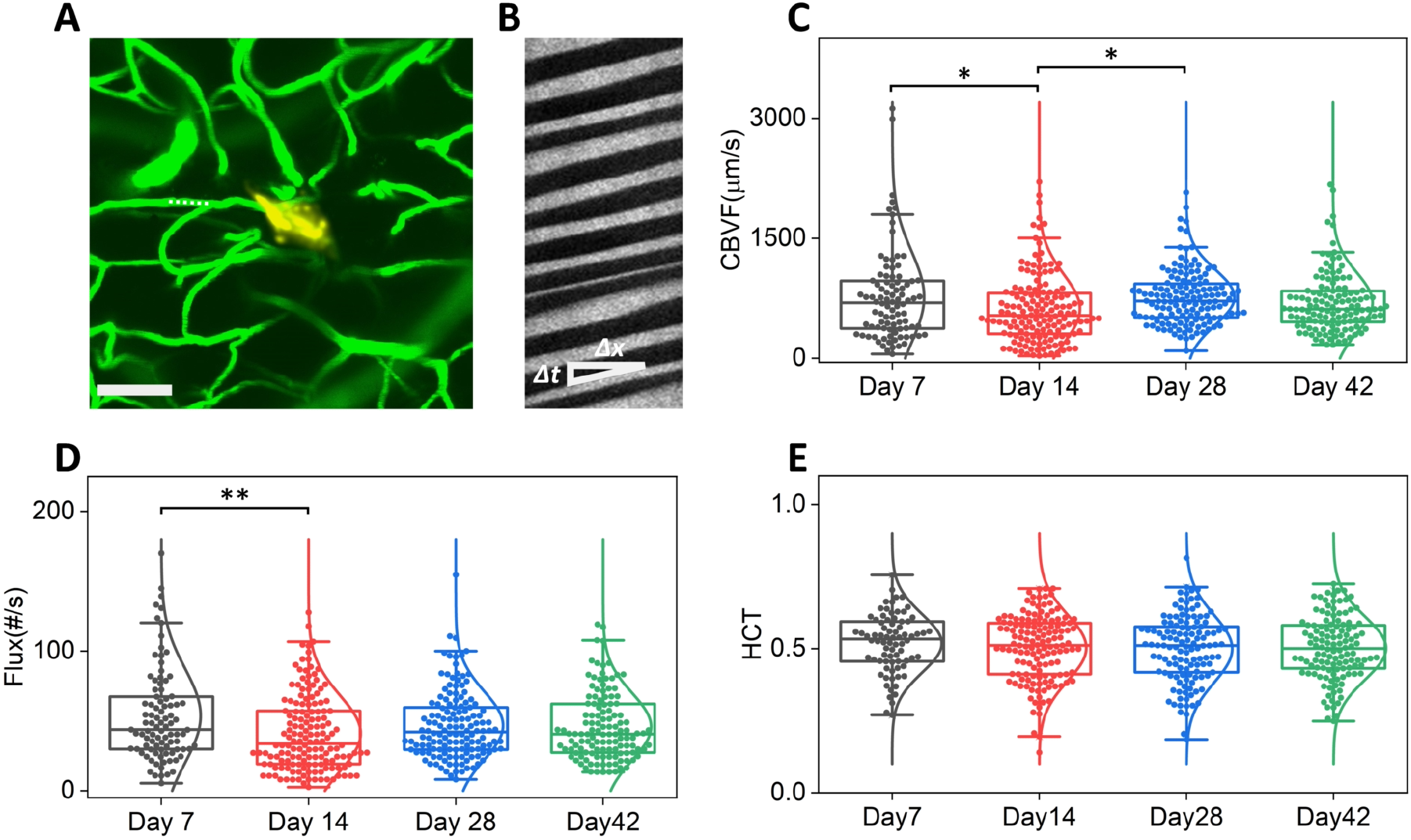
Microcirculation in perfused capillaries near the implantation sites remained stable over time. **A:** representative 2P imaging marking the position of a capillary (dashed line) for the line scans in **B**. Scale bar: 200 μm. Green, vasculature; golden Yellow: NET; **B**: Matrix of line scans showing movement of RBCs as dark stripes, the slope of which gives the blood flow speed. **C** – **E**: Box plot of RBC velocity **(C)**, flux **(D)**, and hematocrit **(E)** values from week 1 to week 6 post implantation. Whisker plots show the 25th percentile and the 75th percentile of the data set. For **(C)** *η*_p_^2^ = 0.022, \**P* < 0.05; *n* = 513 capillaries (6 mice); RBC velocity; **(D)** *η*_p_^2^ = 0.023, \*\**P* < 0.01; *n* = 515 capillaries (6 mice); RBC flux; **(E)** *n* = 466 capillaries (6 mice); hematocrit, by one-way ANOVA.

## 4. Discussion

Using longitudinal imaging of vasculature and microcirculation combined with electrophysiological recordings six weeks following the implantation of ultraflexible NETs, we characterized the degree and duration of the structural and functional consequences in neural and vascular networks associated with the co-implantation of ultraflexible NETs and a cranial window to best use such a preparation in aged animals. Heightened angiogenesis and vascular remodeling in the first two weeks after implantation resulted in restoration of perfused vessels near NETs, during which period the recorded LFPs and unit activities rapidly increased and reached saturation. While vascular remodeling progressed with shallow cortical layers often preceding deeper layers, little difference in layer 2/3 and layer 4/5 neural activity was detected. Occasional vascular abnormalities appeared in this period and subsided later. These results support the application of ultraflexible NETs in aged animals at minimal perturbations to endogenous neurophysiology after a recovery period from surgical implantation depends on the application. To eliminate the influence of implantation trauma, a recovery period of two weeks after NET implantation will suffice for populational-based studies on electrophysiological activity, and a longer period of four to six weeks is necessary for studies with a microscopic vascular component, such as neurovascular interactions. The time course of recovery is comparable with our previous studies using young, adult mice. It is worth noting that the implantation of a cranial window itself induces tissue inflammatory responses such as elevated glial activation and high spine turnover(*43, 44*) that subsides in 2 – 3 weeks, which aligns with the time needed for tissue healing after NET implantation.

We detected variations in the acute implantation damage and its progression. Among factors that may play important roles in the implantation trauma, we did not have a fine control on the implantation speed. We manually operated the micro-manipulator on the stereotaxic instrument and inserted NETs at a speed of about 0.2 – 0.4 mm/s. An optimal insertion speed that balances tissue dimpling at low speeds and rapture of vessels and cells along implantation at high speeds may result in better tissue-NET interface, faster post-implantation recovery, and more promptly restored electrophysiological activity.

We used an additional group of animals implanted with 4-shank, 128-channel NETs to improve single unit detection fidelity and unit counts, so that we could boost depth resolution in recording and perform clustering analysis for putative cell classifications. The number of recorded single units varied significantly with time in the first two weeks. However, the ratio between putative narrow-waveform interneuron and pyramidal cells remained stable. This suggested that even in the most dynamic phase after NET implantation, there were no changes in the proportion of active neuronal types that may otherwise influence the circuits’ ability to generate recurrent excitation or excessive inhibition. This group of animals did not go through longitudinal 2P imaging. Their electrophysiological recording performance over time is similar to the group that underwent extensive imaging, suggesting that longitudinal imaging employed in this study did not induce noticeable phototoxic effect on neural electrophysiological activity.

## Acknowledgments

This work was funded by the National Institute of Neurological Disorders and Stroke, under R01NS109361 (to L.L.), U01 NS115588 (to C.X.), and R01NS102917 (to C.X.).

## Data availability statement

The raw/processed data required to reproduce these findings cannot be shared at this time due to technical or time limitations.

## Supplementary methods, figures, video and table

### Supplementary figures

**Supplementary Figure S1.**
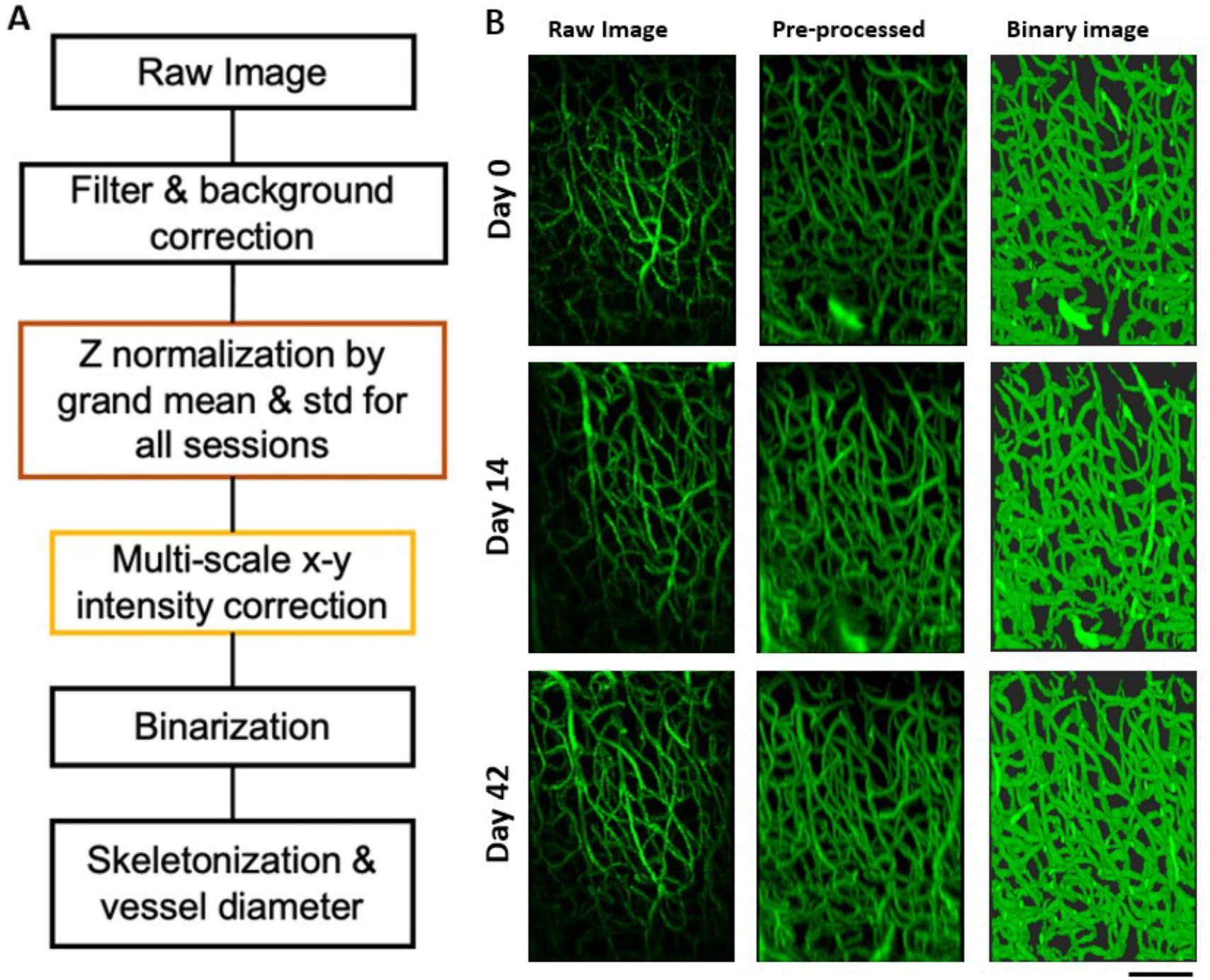
Image processing steps to quantify the vascular recovery. **A:** Flow chart of 2P image processing. Raw images were preprocessed using Gaussian filtering, background correction, z-normalization and x-y normalization. Then they were threshold for vasculature volume fraction analysis and converted to the diameter map to estimate the diameter change across time. **B:** Representative 3D view of raw images, pre-processed images and binary images at different time sessions. Scale bar: 100 μm.

**Supplementary Figure S2.**
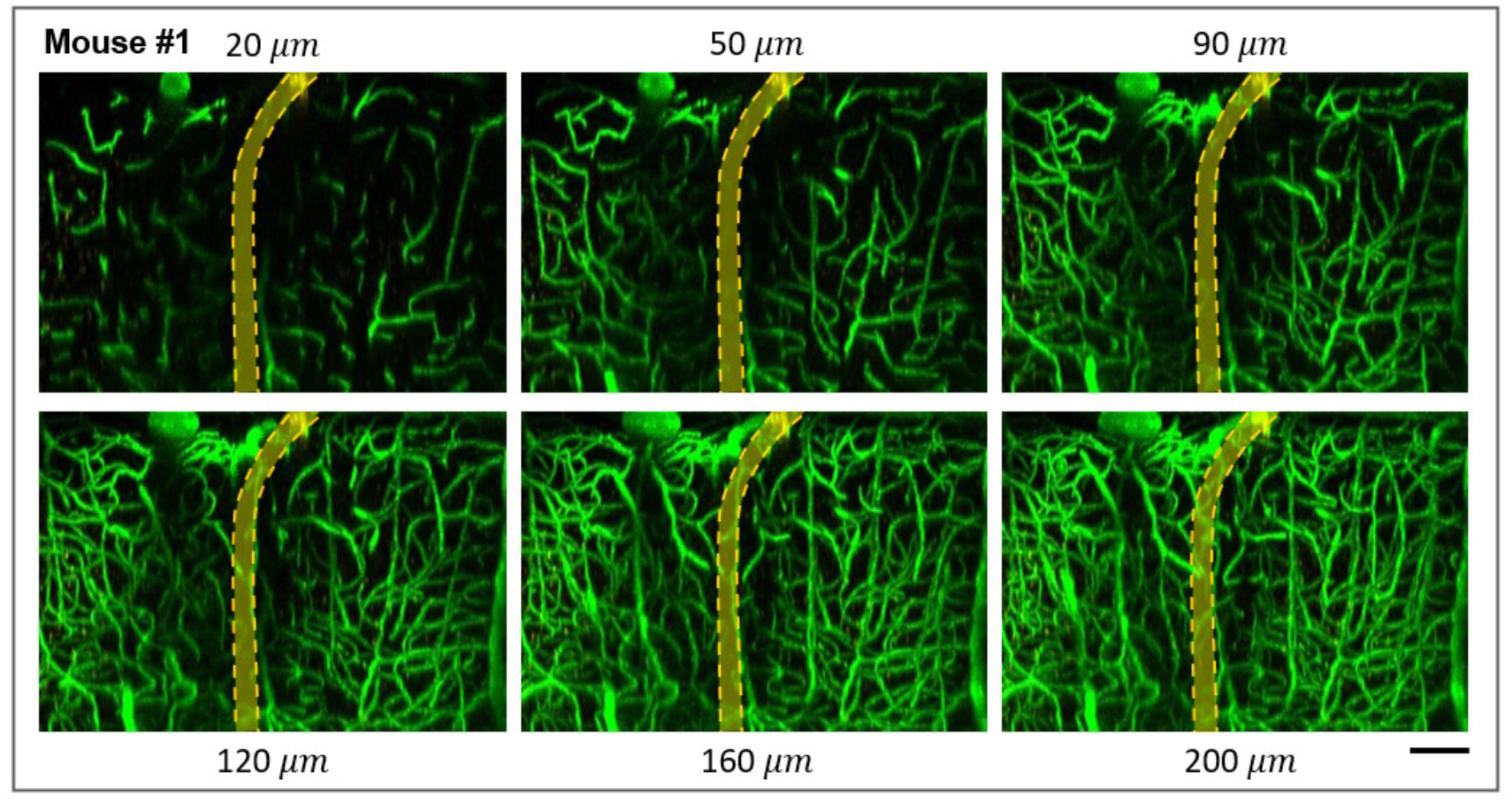
A range of thickness (20μm, 50μm, 90μm, 120μm, 160μm, 200μm,) along the third dimension (y axis) for MIP were tested. 90 μm was chosen to optimize visualization of the microscopic injury and its recovery over time. Scale bars: 100 μm.

**Supplementary Figure S3.**
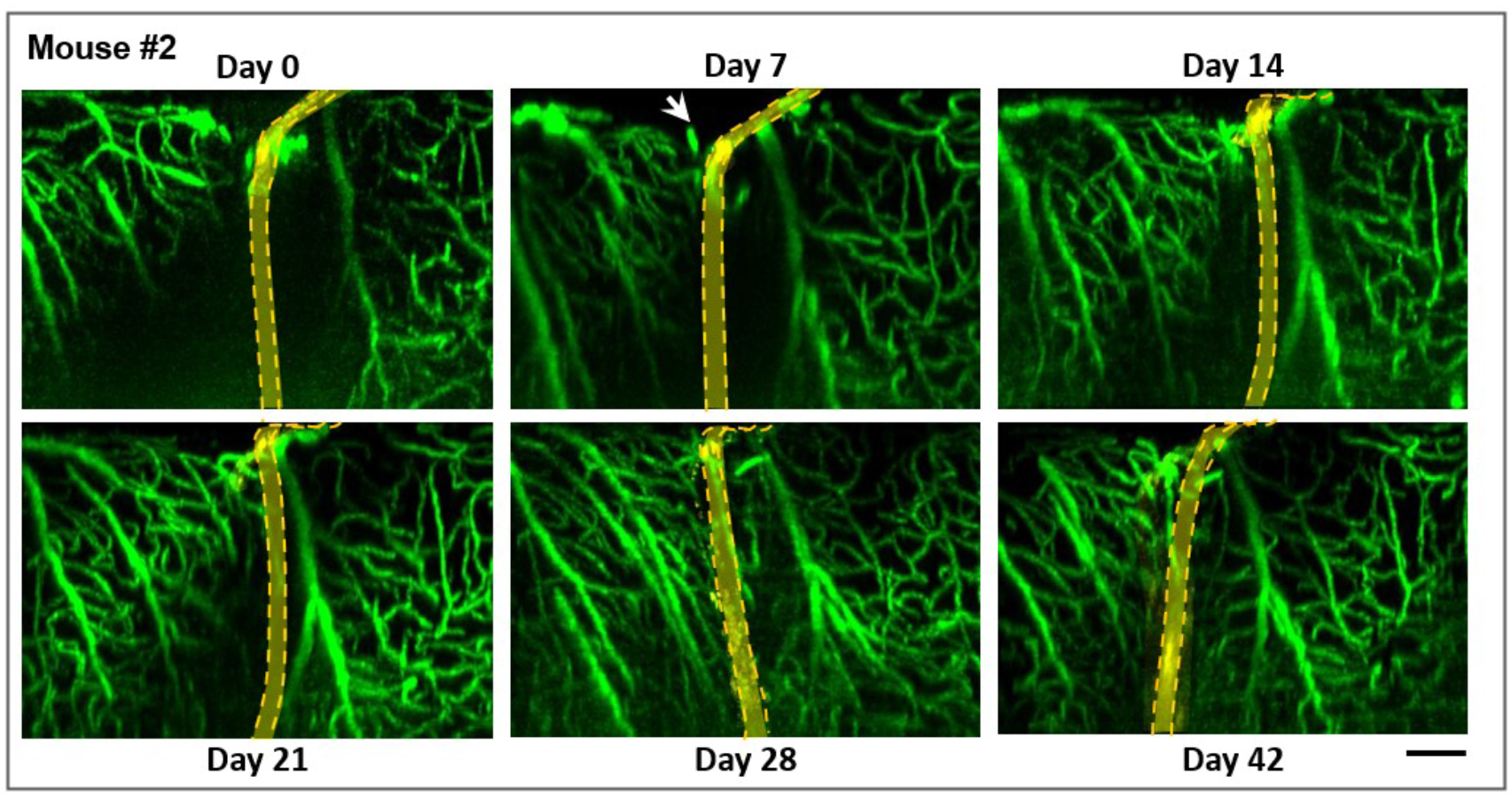
A more severe implantation injury than what is shown in Fig. 2 to demonstrate the variations of the implantation trauma among animals. Arrow marks a ‘dint’, deformation of the brain tissue, at Day 0 and Day 7 after the implantation. Scale bar: 100 μm.

**Supplementary Figure S4.**
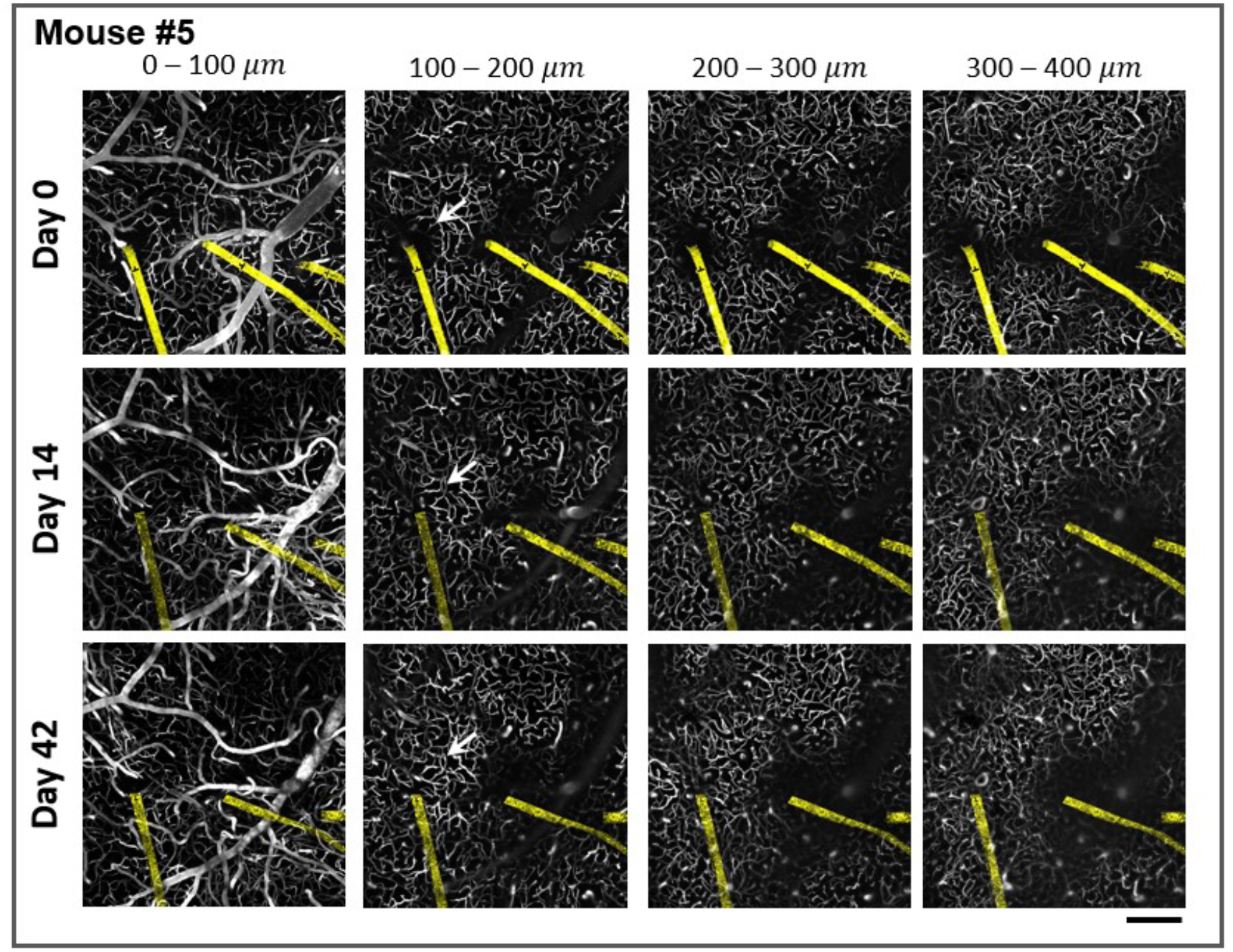
Variations in the vascular recovery at different locations in the same animal. Arrow shows an implantation site away from major pial vessels had faster vascular recovery than the other two sites. Scale bar: 200 μm.

